# A small-molecule lycorine derivative protects from obesity by targeting Na^+^/K^+^-ATPase *α*3 independent of food intake suppression

**DOI:** 10.1101/2022.08.24.505199

**Authors:** Henan Zhang, Yunfu Zhou, Duozhi Chen, Yibao Zhang, Fei Jin, Zhangcheng Chen, Chen Li, Boran Chang, Rong Zeng, Jinsong Li, Xiaomin Song, Yan Chen, Xiaojiang Hao, Lin Li

**Author notes:** Corresponding author. Lin Li and Xiaojiang Hao. These authors contributed equally to this work (Henan Zhang, Yunfu Zhou, Duozhi Chen).

## Abstract

Obesity remains a severe global public health challenge, with contemporary therapeutic approaches primarily focusing on appetite suppression. Here, we discovered HLY72, a lycorine-derived small molecule that potently counteracts obesity in mice at doses without affecting food intake, by promoting sympathetic activation-mediated lipolysis and thermogenesis in adipose tissues. Mass spectrometry identified Na^+^/K^+^-ATPase (NKA) α3, the brain-specific isoform, as HLY72’s target. Only blood-brain barrier-permeable NKA inhibitors reproduce HLY72’s anti-obesity effects, and resistance to HLY72 treatment occurs specifically in NKA α3 (not α1) knockin mice harboring a HLY72-binding mutation. Similar to the chemical inhibition by HLY72, genetic inhibition of NKA α3 also effectively protects mice from diet-induced obesity. These findings point NKA α3 as a potent anti-obesity drug target and highlight HLY72’s potential in treating and preventing obesity independent of appetite control.

## INTRODUCTION

Control of excess body fat is one of the greatest healthcare challenges of our time. In 2021, the global prevalence of obesity had increased by 155.1% in males and 104.9% in females compared with 1990, and China had the largest population of adults with overweight or obesity (402 million individuals) (*1*). Obesity has been recognized as a significant risk factor for a variety of serious conditions such as type 2 diabetes, atherosclerosis, cardiovascular disease, hypertension, and even cancers (*2, 3*). Over the past decades, great efforts have been made to explore obesity pathophysiology and develop anti-obesity medications (*4–11*). Currently available anti-obesity medications largely act on central appetite pathways to reduce caloric consumption (*12*). For example, glucagon-like peptide-1 receptor (GLP1R) agonists, the most successful weight-loss therapeutics so far, affect body weight mainly through regulating food intake. (*13, 14*). Of note, drugs acting by suppression of food intake generally cause related adverse reactions such as nausea, vomiting, depression, and headache (*15*). Therefore, it would be of great value to develop therapeutics that could protect from obesity without the need to change feeding patterns.

Na^+^/K^+^-ATPase (NKA) is a ubiquitous membrane transport protein, and is responsible for maintaining Na^+^ and K^+^ gradients across the cell membrane, thereby influencing cell volume, absorption, secretion, excitability, and body homeostasis (*16*). NKA contains two key functional subunits, α and β subunits. The α subunit is the catalytic unit and the target for pharmacological intervention, while the β subunit plays a regulatory role mainly through facilitating the correct folding, membrane localization, and stability of the α subunit (*17, 18*). In mammals, there are four α isoforms (α1-4) of NKA, which are expressed differentially in tissues and have distinct functions. α1 isoform is ubiquitously expressed and may play a “housekeeping” role, while the rest three α isoforms of NKA have a more specific tissue distribution−α2 mostly in skeletal muscle and heart; α3 in neurons; and α4 predominantly in testes (*19*). In addition to its activity as an electrogenic pump, NKA α1 also acts as an oxidant signaling transducer, which modulates reactive oxygen species (ROS) amplification through an interaction with Src kinase (*20*). It was reported that a peptide targeting NKA α1-Src interaction could alleviate adiposity (*21*). In the current study, starting with a small-molecule compound HLY72 as a chemical probe, we uncovered NKA α3 as a promising drug target to treat and prevent obesity. Our data showed that either chemical or genetic inhibition of NKA α3 protects mice from obesity and related defects. Importantly, our data showed that HLY72 could potently counteract obesity and ameliorate metabolic impairments at doses without altering food intake.

## RESULTS

### Lycorine derivative HLY72 elicits weight loss in mice

In our previous work, we identified the small-molecule lycorine derivative HLY78 as a Wnt agonist, with its analog HLY72 serving as a negative control (**Fig. 1A**) (*22*). When testing the possible effects of HLY78 on mice, we found that intraperitoneal administration of both HLY78 and HLY72 at 10 mg/kg resulted in a significant reduction in body weight of mice fed a normal chow diet (NCD) (**Fig. 1, B and C**), suggesting that this weight-loss activity is independent of Wnt/β-catenin signaling. This unexpected finding caught our attention. To rule out potential Wnt pathway activation effects, we used the Wnt-inactive analog HLY72 to further investigate the HLY78/HLY72-induced weight-loss phenotype. First, we tested the effect of HLY72 at varying doses on dark phase food intake in mice fed a NCD. The result showed that HLY72 did not affect food consumption at doses ≤ 10 mg/kg, while at doses higher than 15 mg/kg, HLY72 began to inhibit food intake (**Fig. 1D**). Next, we examined the effect of HLY72 on body weight of obese mice fed a 60% high-fat diet (HFD) at varying doses ≤10 mg/kg (**Fig. 1E**). The result showed that HLY72 significantly impeded body weight gain in a dose-dependent manner across all tested doses (**Fig. 1F**). Measurement of cumulative food intake showed that none of these doses significantly affected food intake in mice (**fig. S1A**). These results indicate that the anti-obesity effect of HLY72 is independent of appetite suppression at least within a certain dose range. Therefore, to exclude food intake suppression as a contributing factor, we used HLY72 at a dose of 10 mg/kg unless otherwise specified in subsequent studies. Compared to the vehicle group, HLY72 treatment significantly reduced body fat without affecting lean mass (**Fig. 1, G and H, and fig. S1, B and C)**. We then conducted metabolic cage studies on the mice treated with HLY72 or vehicles for 3 weeks (**Fig. 1E)**. Compared to the vehicle control, HLY72 treatment significantly increased oxygen consumption (VO_2_) and energy expenditure (EE) during the dark cycle, with no change in physical activity (**Fig. 1, I to K**). Of note, HLY72 treatment resulted in a decreased respiratory exchange ratio (RER) as compared to the vehicle control (**Fig. 1L),** indicating an enhanced lipid oxidation. Next, we performed the glucose tolerance test (GTT) and insulin tolerance test (ITT) on the mice treated with HLY72 for 6 weeks (**Fig. 1E**). Glucose clearance from the circulation during GTT was significantly faster in HLY72-treated mice than in the vehicle-treated control mice (**Fig. 1M**). Meanwhile, insulin sensitivity in HLY72-treated mice was also dramatically improved as reflected by ITT (**Fig. 1N**). Moreover, HFD-induced hepatic steatosis was ameliorated in HLY72-treated mice (**Fig. 1O**). Together, these results showed that HLY72 could effectively protect mice from HFD-induced obesity and related metabolic dysfunctions without affecting food intake.

**Figure 1.**
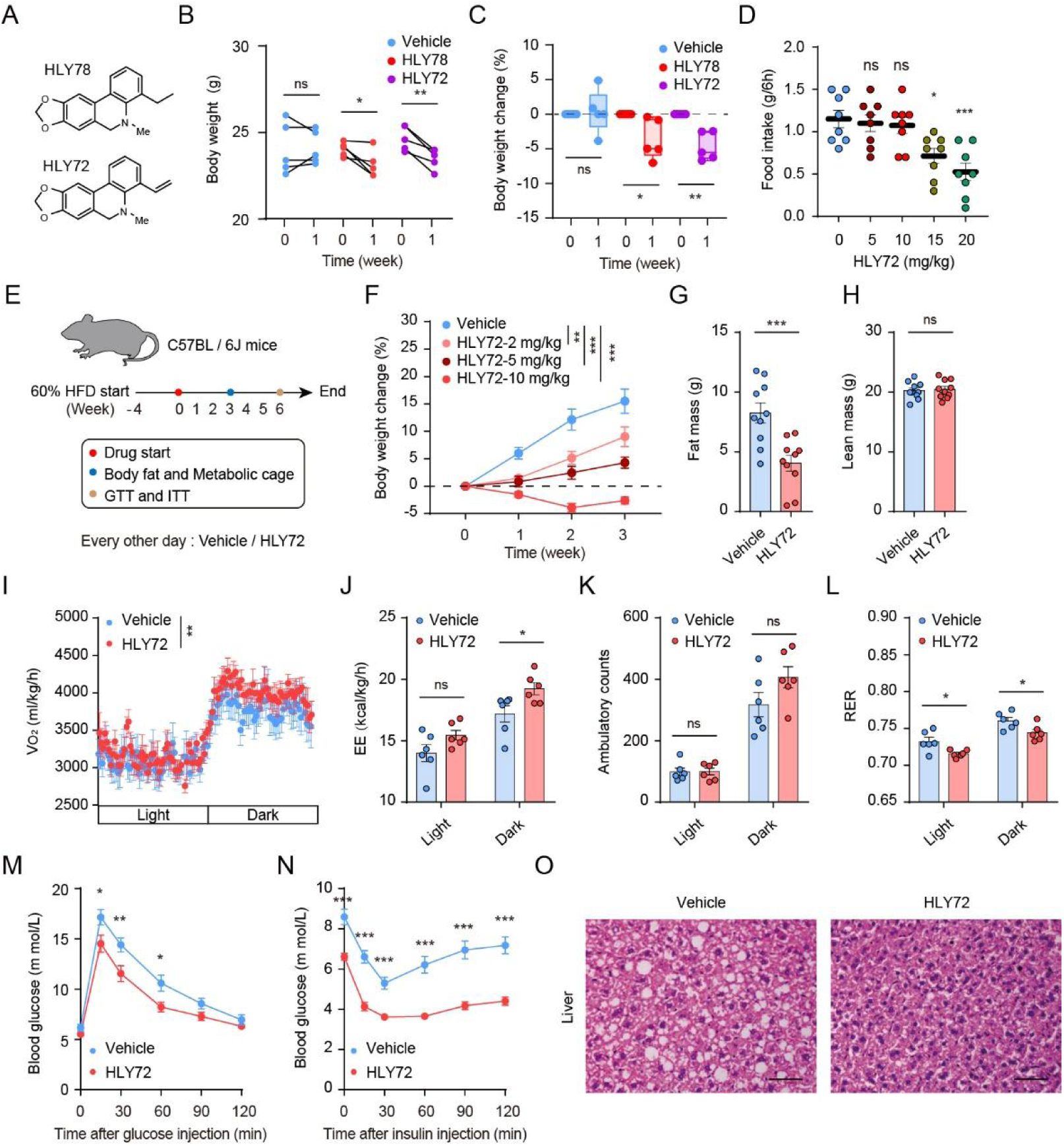
HLY72 protects mice from obesity and metabolic dysfunction without affecting food intake. (**A**) Chemical structure of HLY78 and HLY72. (**B** and **C**) Body weight (B) and body weight change (C) of 10-week-old male mice treated with vehicle, HLY78 or HLY72 at a dose of 10 mg/kg. n=5. (**D**) Dose-response analysis of HLY72 on food intake of 8-week-old male mice. n=8. (**E**) Experimental scheme for (G to R) on 8-week-old male mice. (**F**) Dose-dependent effects of HLY72 on the body weight change. n=10. Relevant food intake information is shown in Fig. S1A. (**G** and **H**) Fat mass (G), lean mass (H). n=10. HLY72: 10 mg/kg. (**I** to **L**) VO_2_ (I), EE (J), ambulatory counts (K) and RER (L). n=6. HLY72: 10 mg/kg. (**M** to **O**) GTT (M), ITT (N), and HE staining of liver sections (O). n=9. HLY72: 10 mg/kg. Scale bar, 50 mm. Data are mean ± s.e.m. *p* values were determined by two-tailed paired student’s test in [(B), (C)], two-tailed un-paired student’s test in [(G), (H)], one-way ANOVA in [(D)] and two-way ANOVA in [(F), (I) to (N)]. **\****p*<0.05, **\*\****p*<0.01, **\*\*\****p*<0.001 compared to control. ns, not significant with *p*>0.05. Data points represent individual mice [(B) to (D), (G), (H), and (J) to (L)].

### HLY72 stimulates both lipolysis and thermogenesis in adipose tissues

Adipocytes are crucial for maintaining energy homeostasis. To explore the mechanisms underlying the weight-loss effect of HLY72, we first assessed its impact on lipolysis in adipocytes. Hormone-sensitive lipase (HSL) is a key enzyme catalyzing lipolysis in the white adipose tissue and its activity is reflected by its phosphorylation; while, protein kinase A (PKA) functions upstream of p-HSL and its activity is reflected by protein levels of phosphorylated PKA substrates (*23, 24*). After intraperitoneal administration of HLY72 in 8-week-old mice for 6 hours, we observed a significant increase in the levels of p-HSL and p-PKA substrates in epididymal white adipose tissue (eWAT). (**Fig. 2, A and B**). Consistent with these results, glycerol and free fatty acid (FFA) were increased in eWAT after HLY72 treatment (**Fig. 2, C and D**). The sympathetic nervous system (SNS) plays a key role in regulating lipolysis. Sympathetic activation leads to the elevation of norepinephrine (NE), which binds to β-adrenergic receptors of white adipose tissue and consequently stimulates lipolysis (*25*). To test whether HLY72 might affect lipolysis through SNS, we measured NE in eWAT and serum of mice under HLY72 or vehicle treatment. Compared to the vehicle control, HLY72 treatment induced an increase of NE in both eWAT and serum (**Fig. 2, E and F**), suggesting that HLY72 might regulate lipolysis through sympathetic activation. To validate this, we introduced hexamethonium, a sympathetic ganglion blocker (*25*), and results showed that the increased phosphorylation of HSL induced by HLY72 could be significantly blocked by hexamethonium (**Fig. 2, G and H**), supporting that HLY72 functions mainly through an activation of the sympathetic system. Sympathetic activation generally could stimulate both lipolysis and thermogenesis (*26*). In addition, we observed an increased oxygen consumption elicited by HLY72 treatment in our metabolic cage studies above (**Fig. 1I**), which also implied a function of HLY72 in thermogenesis. To investigate whether HLY72 could affect thermogenesis, we intraperitoneally administered HLY72 or vehicle to mice housed at room temperature (22°C), then exposed them to a cold environment (4°C) for 6 hours. Compared with the vehicle group, the rectal temperature of HLY72-treated mice was significantly increased (**Fig. 2I**). Consistent with this, HLY72 treatment significantly upregulated mRNA levels of thermogenic genes, including *UCP1*, *Cieda, PGC1α,* and *Dio2* in BAT, as reflected by the quantitative PCR analysis (**Fig. 2, J to M**). Meanwhile, UCP1 protein expression in BAT was also significantly increased as reflected by Western blot analysis (**Fig. 2, N and O)**. Collectively, these results suggest that HLY72 promotes both lipolysis and thermogenesis in adipose tissues, likely through activation of the sympathetic nervous system.

**Figure 2.**
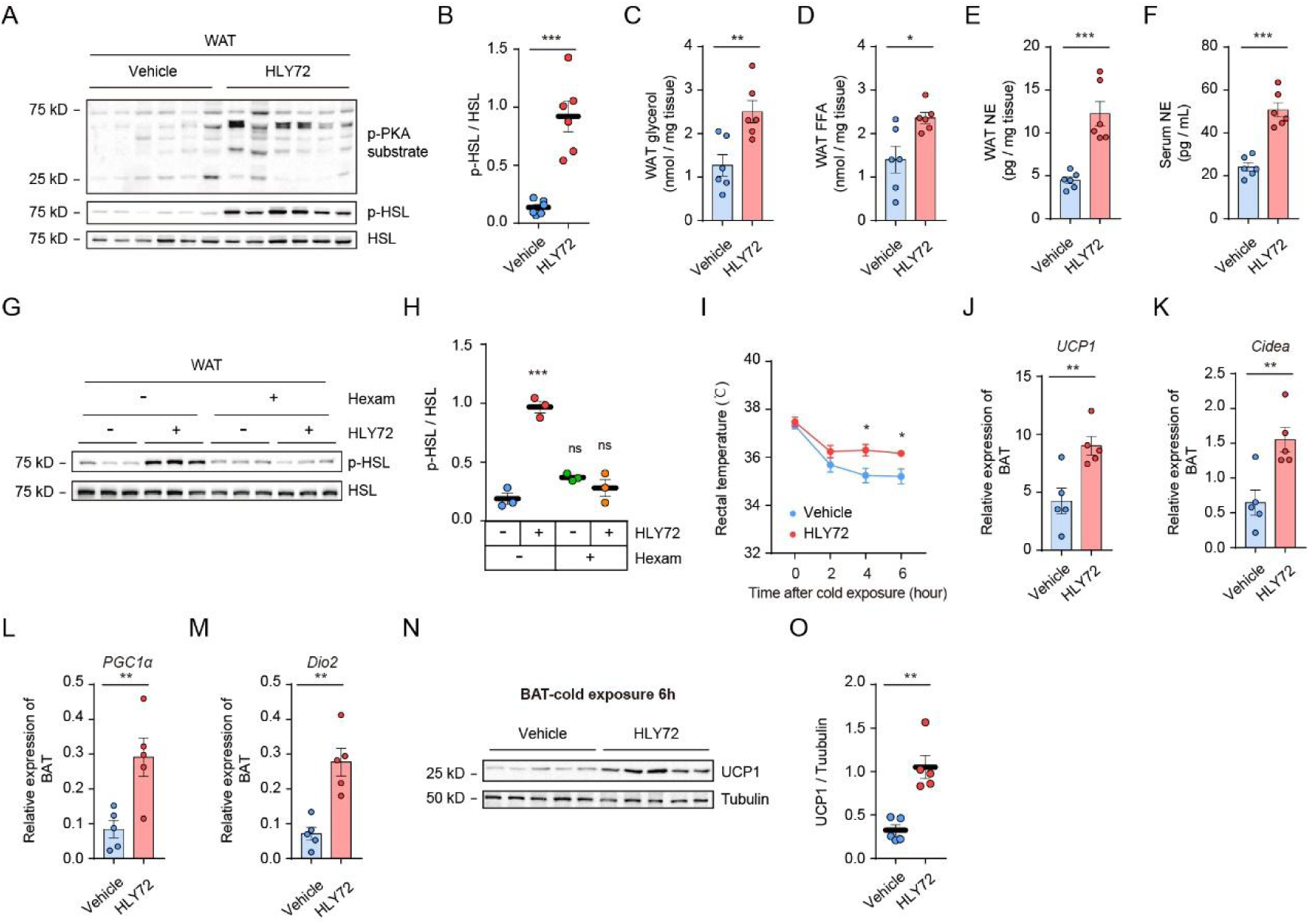
HLY72 promotes WAT lipolysis and BAT thermogenesis mainly through increasing sympathetic activity. 8-week-old mice were i.p. injected with 10 mg/kg of HLY72 or vehicle for 6 hours. (**A** and **B**) p-HSL and p-PKA substrates in total protein extracts of epididymal fats were examined by immunoblot analysis (A) and analyzed statistically (B). n=6. (**C** and **D**) Glycerol (C) and FFA (D) of epididymal fats were analyzed by ELISA. n=6. (**E** and **F**) Epididymal fats (E) and serum (F) levels of norepinehrine (NE) were analyzed by ELISA. n=6. (**G** and **H**) Mice were treated with HLY72 in combination with 20 mg/kg hexamethonium. p-HSL levels in total protein extracts of epididymal fats were examined by immunoblot analysis (G), and analyzed statistically (H). n=3. (**I**) Rectal temperatures of vehicle- and HLY72-treated mice during cold exposure (4 ℃) for 6 hours. n=5. (**J** to **M**) Transcriptional levels of thermogenesis-relevant genes in BAT from mice treated with HLY72 or vehicle after cold exposure (4 ℃). n=5. (**N** and **O**) UCP1 protein levels in BAT after cold exposure (4 ℃) were examined by immunoblot analysis (N) and analyzed statistically (O). n=5. Data are mean ± s.e.m. *p* values were determined by two-tailed un-paired student’s test in [(B) to (F), (J) to (M) and (O)], one-way ANOVA in [(H)] and two-way ANOVA in [(I)]. \**p*<0.05, \*\**p*<0.01, \*\*\**p*<0.001. ns, not significant with *p*>0.05. Data points represent individual mice [(B) to (F), (H), (J) to (M), and (O)].

### Na^+^/K^+^-ATPase *α3* is the cellular target of HLY72

To identify the potential cellular target of HLY72, we synthesized biotinylated HLY72 (Bio72) (**fig. S2**) and verified its activity in increasing p-HSL levels, which was comparable to that of HLY72 (**fig. S3, A and B**). Given the fact that the brain is considered the center of energy homeostasis regulation (*27, 28*), we then performed mass spectrometry (MS) analysis for proteins pulled down by Bio72 from mouse brain lysates (**fig. S3C**). Our MS analysis yielded 6 candidates captured only by Bio72 but not by the biotin control (**table. S1**). Among the 6 candidates, NKA α3 attracted our attention due to the reported effects of NKA α1 isoform on body weight (*21*). Consistent with previous reports (*19*), our gene expression analysis showed that NKA α3 was specifically distributed in the brain (**fig. S3D**).

To test if NKA α3 is a potential target of HLY72, we examined the kinetics of possible NKA-HLY72 interactions by using two distinct biophysical techniques—surface plasmon resonance (SPR) and microscale thermophoresis (MST). Due to a relatively large amount of protein samples required for these two assays and the high homology between different isoforms of NKA, we purified NKA α1 from pig kidney using a method described by Klodos *et al*. (*29*) (**Fig. 3A**). These two methods produced similar results with the dissociation constant (K_D_) for NKA-HLY72 interaction being 4.2 and 3.2 μM respectively (**Fig. 3B and fig. S3E**). Next, we examined the possible effect of HLY72 on the enzymatic activity of NKA α3. We overexpressed Flag-tagged NKA composed of α3 and β1 subunit in HEK293T cells and purified it by using the Flag affinity chromatography (**Fig. 3C**). We first confirmed that the NKA sample we obtained was active by testing it with the classic NKA inhibitor digitoxin (**fig. S3F**). We then evaluated the inhibitory activity of HLY72 against NKA α3. Our result showed that HLY72 inhibited NKA activity by approximately 40% at a concentration of 10 μM (**Fig. 3D**); and the half-maximal inhibitory concentration (K_i_) of HLY72 was calculated to be 18.6 μM (**Fig. 3E**).

**Figure 3.**
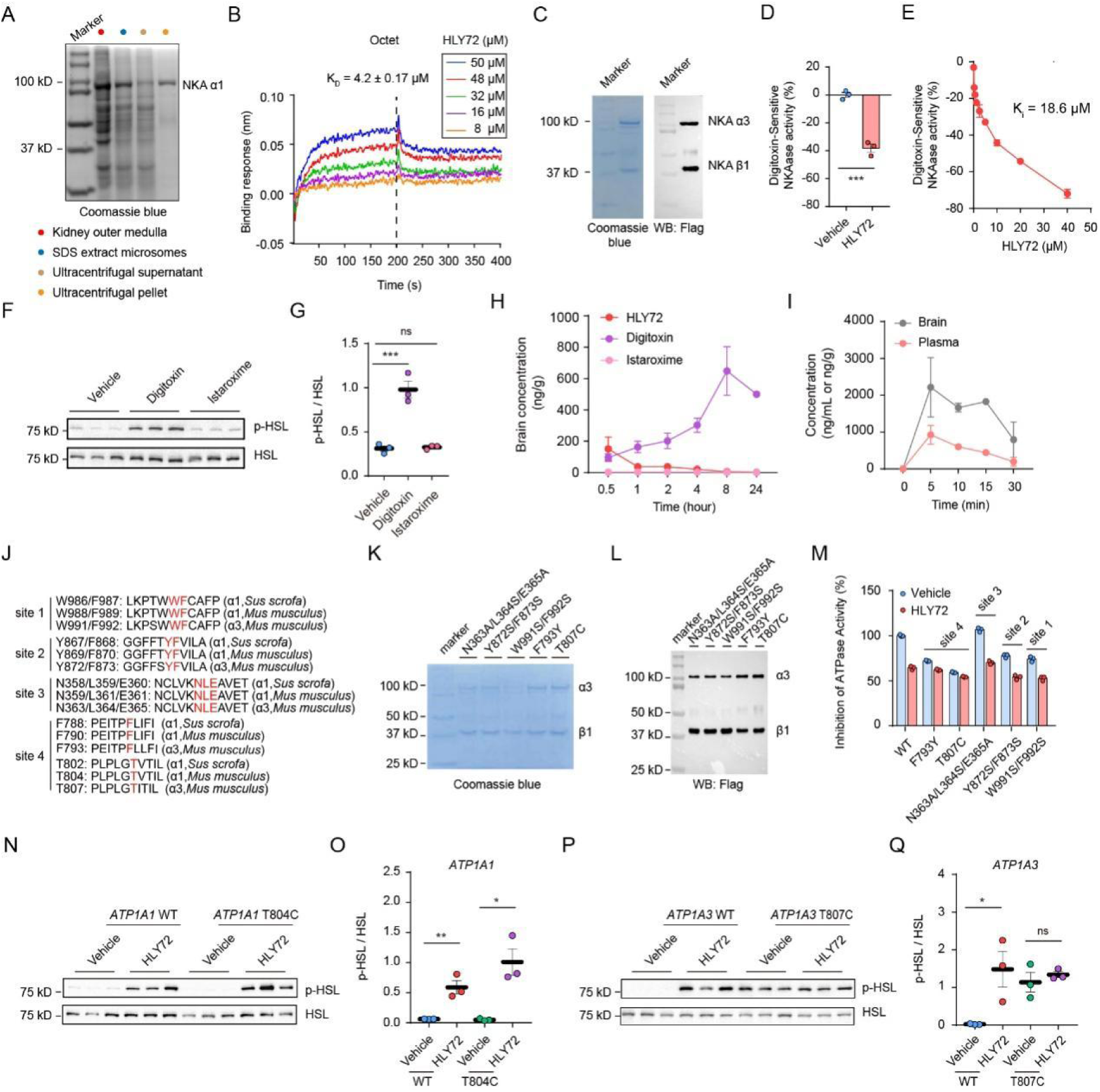
NKA α3 is identified as a target for HLY72. (**A**) The pig kidney outer medulla NKA α1 was purified following the protocol described in the Method. (**B**) Binding affinity of HLY72 with NKA were determined by the biolayer interferometry assay. (**C**) Mouse NKA α3β1 overexpressed in HEK293T was purified. (**D**) The effect of HLY72 (10 mM) on NKA α3β1 activity. (**E**) Dose-dependent inhibition of NKA α3β1 activity by HLY72. (**F** and **G**) p-HSL levels in total protein extracts of epididymal fats from 8-week-old male mice treated with 5 mg/kg of digitoxin or istaroxime were examined by immunoblot analysis (F) and analyzed statistically (G). n=3. (**H**) Time-dependent brain distribution profiles of HLY72, digitoxin, and istaroxime following oral administration at a dose of 3 mg/kg. (**I**) Brain and plasma distribution of HLY72 following intraperitoneal administration at a dose of 10 mg/kg. (**J**) Comparison of amino acid sequences of pig NKA α1 with mouse NKA α1 and α3. (**K** and **L**) Purification of various α3 mutants that were designed based on docking results. (**M**) Evaluation of NKA α3 mutants for their sensitivities to HLY72 treatment. (**N** to **Q**) p-HSL levels for *ATP1A1* T804C mice (N and O) and *ATP1A3* T807C 8-week-old male mice (P and Q) treated with 10 mg/kg HLY72 for 6 hours were examined by immunoblot analysis (N and P) and analyzed statistically (O and Q). n=3. Data are mean ± s.e.m. *p* values were determined by two-tailed un-paired student’s test in [(D), (O) and (Q)], one-way ANOVA in [(G)]. **\****p*<0.05, **\*\****p*<0.01, **\*\*\****p*<0.001. ns, not significant with *p*>0.05. Data points represent individual mice [(G), (O) and (Q)].

If HLY72 executes its anti-obesity activity by binding and inhibiting NKA, other NKA inhibitors are supposed to have similar effects as HLY72. To test this possibility, we introduced two cardiac glycosides (CGs), digitoxin and istaroxime, and we confirmed the inhibitory activity of istaroxime against NKA α3 in our system (**fig. S3G**). Although both compounds exhibited inhibitory activity against NKA *in vitro*, only digitoxin—but not istaroxime—significantly increased the phosphorylation of HSL in eWAT (**Fig. 3, F and G**). Digitoxin is reported to be capable of going through the blood-brain barrier, but there is no such information available for istaroxime (*30, 31*). If the brain-localized NKA α3 was responsible for the effects we observed above, it is reasonable to assume that NKA inhibitors function as anti-obesity agents only when they can access the brain. To test this, we examined the brain distribution of HLY72, digitoxin, and istaroxime. As expected, HLY72 and digitoxin, but not istaroxime, could be detected in the brain; moreover, HLY72 exhibited transient brain retention compared to the higher stability of digitoxin **(Fig. 3H)**. Further metabolic characterization revealed that HLY72 exhibited a short half-life of only several minutes (**Fig. 3I**). This result further supports that the NKA α3, which is dominantly distributed in the brain, could be the target for HLY72. Given that the half-life of digitoxin is approximately 7 days (*30*), we evaluated digitoxin’s weight-reduction effects using the same protocol as for HLY72 except with a modified dosing interval of once every 7 days (**fig. S3H)**. Digitoxin exhibited significant anti-obesity efficacy when administered at a dose of 7.5 μg/kg (**fig. S3I)** without affecting food intake (**fig. S3J**). Moreover, digitoxin delivered at 7.5 μg/kg led to reduced body fat as compared with the controls (**fig. S3, L and M**). Meanwhile, digitoxin treatment also significantly upregulated UCP1 protein expression in BAT (**fig. S3, N and O**).

Based on the crystal structure of pig NKA α1 complexed with digoxin (Protein Data Bank 7DDI) (*31, 32*), we predicted possible binding sites for HLY72 on NKA by using the AutoDock Vina docking program (*33*). Four possible binding cavities were identified on the pig NKA α1 subunit surface, and all of them are conserved in mouse α3 (**Fig. 3J**). Among these four sites, site 4, encompassing T807 and F793, is also the classic binding site of NKA for CGs, including digitoxin (*34, 35*). Next, we generated corresponding mutants for these four sites on mouse α3 and purified them (**Fig. 3, K and L**). We then performed the NKA activity assay, and the result showed that mutations targeting site 4, especially T807C, almost eliminated HLY72’s inhibitory activity against NKA (**Fig. 3M**). This result implied that HLY72 binds to the same site on NKA as digitoxin. Of note, mutations of sites 1, 2, and 4 all decreased NKA’s ATPase activity, with the T807C mutation resulting in a decrease of ∼50%. This is consistent with previous reports that the T807C mutation decreases NKA’s ATPase activity (*36*).

To investigate whether NKA α3 is the bona fide endogenous target for HLY72, we tried to construct *ATP1A1* T804C, *ATP1A2* T801C, and *ATP1A3* T807C point mutation knockin mice through the zygote injection method (*37*). We successfully generated homozygous *ATP1A1* T804C and *ATP1A3* T807C mice (**fig. S4, A and B**), but the generation of *ATP1A2* T801C mice failed. We first examined the impact of high-dose digitoxin and HLY72 on food intake in *ATP1A1* T804C and *ATP1A3* T807C mice. After injection of digitoxin at 150 μg/kg or HLY72 at 20 mg/kg, the food intake of *ATP1A1* T804C mice was similarly reduced compared to WT mice; while neither digitoxin nor HLY72 could induce any apparent effect on food intake in *ATP1A3* T807C mice (**fig. 4, C and D**). Moreover, either HLY72 or digitoxin treatment increased p-HSL levels in eWAT of *ATP1A1* T804C mice to a degree comparable to WT mice with basal p-HSL levels remaining unchanged (**Fig. 3, N and O, and fig. S4, E and F**). By contrast, *ATP1A3* T807C mice exhibited significantly elevated basal p-HSL levels compared to *ATP1A3* WT mice, supporting a specific role of NKA α3 in lipolysis regulation; meanwhile, neither HLY72 nor digitoxin treatment could further increase p-HSL levels in *ATP1A3* T807C mice **(Fig. 3, P and Q, and fig. S4, G and H)**. Together, these results showed that the mutation in NKA α3, but not in α1, prevents mice from responding to HLY72/digitoxin treatment, validating NKA α3 as the bona fide target for HLY72.

**Figure 4.**
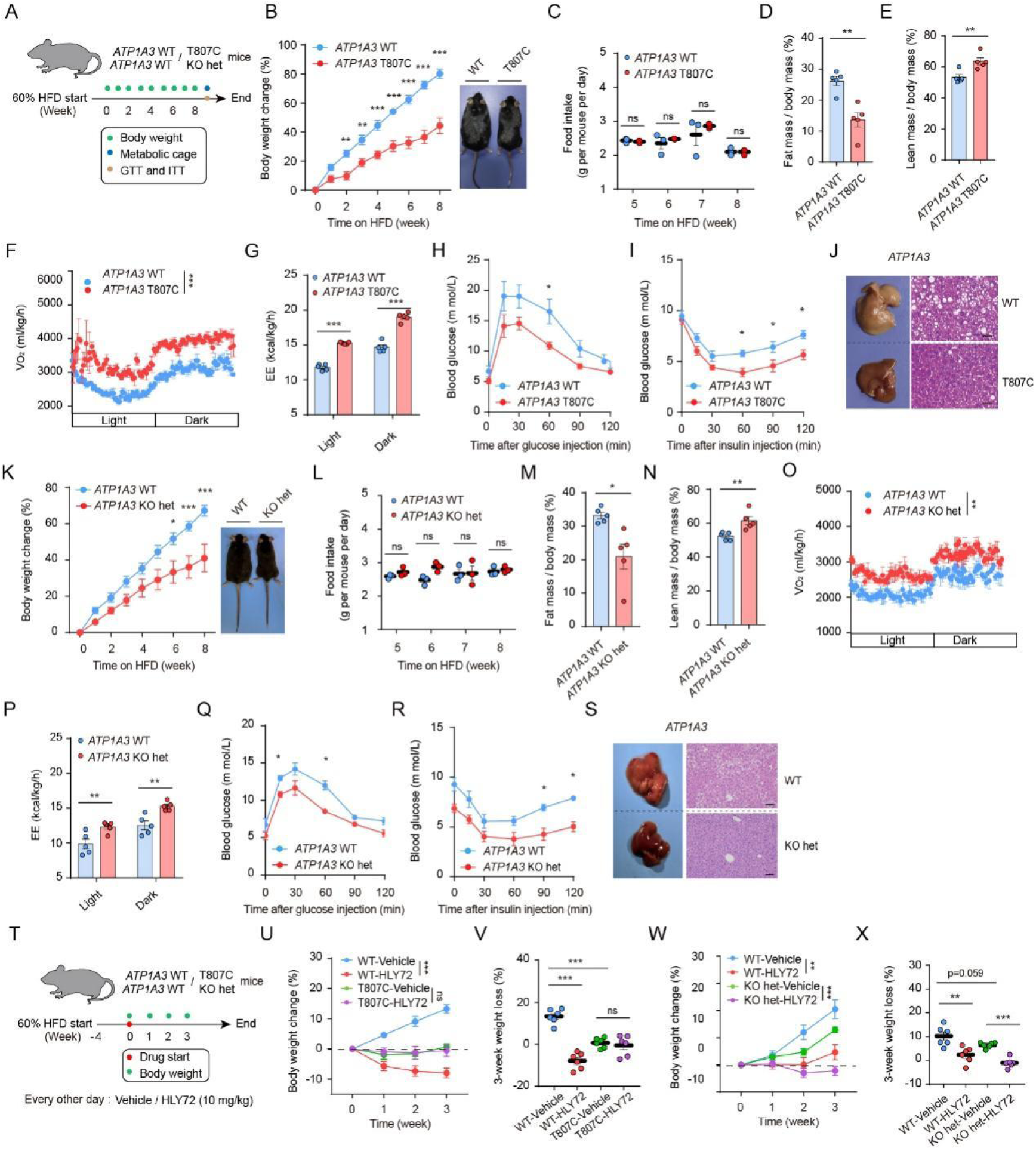
Genetic inhibition of NKA α3 protects mice from HFD-induced obesity and metabolic defects. (**A**) Experimental scheme for (B to S). 8-week-old male mice were used in these experiments. (**B** to **E**) Changes of body weight (B), food intake (C), percentage of fat mass (D), and percentage of lean mass (E) of *ATP1A3* T807C *vs* WT mice. n=5. (**F** and **G**) VO_2_ (F) and EE (G) for *ATP1A3* T807C *vs* WT mice. n=5. (**H** and **I**) GTT (H) and ITT (I) for *ATP1A3* T807C *vs* WT mice. n=5. (**J**) HE staining of liver sections of *ATP1A3* T807C *vs* WT mice. Scale bar, 50 mm. (**K** to **N**) Changes of body weight (J), food intake (L), percentage of fat mass (M), and percentage of lean mass (N) of *ATP1A3* KO het *vs* WT mice. n=5. (**O** and **P**) VO_2_ (O) and EE (PN) for *ATP1A3* KO het *vs* WT mice. n=5. (**Q** and **R**) GTT (Q) and ITT (R) for *ATP1A3* KO het mice *vs* WT. n=5. (**S**) HE staining of liver sections of *ATP1A3* KO het mice *vs* WT. Scale bar, 50 mm. (**T**) Experimental scheme for (U to X). 8-week-old male mice were used. (**U** and **V**) Body weight changes of *ATP1A3* T807C and WT mice treated with vehicle or HLY72 during 3 weeks (U) and at the end of the third week (V). n=6. (**W** and **X**) Body weight changes of *ATP1A3* KO het and WT mice treated with vehicle or HLY72 during 3 weeks (W) and at the end of the third week (X). n=6. Data are mean ± s.e.m. *p* values were determined by two-tailed un-paired student’s test in [(D), (E), (M), (N), (V) and (X)], two-way ANOVA in [(B), (C), (F) to (I), (K), (L), (O) to (R), (U) and (W)]. **\****p*<0.05, **\*\****p*<0.01, **\*\*\****p*<0.001. ns, not significant with *p*>0.05. Data points represent individual mice [(D), (E), (G), (M), (N), (P), (V) and (X)].

### Genetic inhibition of NKA *α3* protects mice from HFD-induced obesity

Our findings above showed that the chemical inhibition of NKA α3 by HLY72 could effectively protect mice from HFD-induced obesity. Next, we set out to investigate whether genetic inhibition of NKA α3 could have similar effects. For this, we took advantage of the property of the NKA α3 T807C mutation—decreasing NKA activity by ∼50% (**Fig. 3M**), and investigated potential effects of such a genetic inhibition of NKA α3 on metabolic features and lipolysis of HFD-fed mice. On the other hand, we also generated *ATP1A3* KO heterozygous (*ATP1A3* KO het) mice (**fig. S5, A and B)**, which also maintain 50% NKA activity. We treated *ATP1A3* T807C or *ATP1A3* KO het mice with a 60 kcal% HFD for 8 weeks, and compared them with the WT control mice in several metabolic-related parameters as indicated in **Fig. 4A**. For the *ATP1A3* T807C mice, their body weight gains were significantly decreased compared to their WT littermates (**Fig. 4B)**. Food intake was evaluated after one month of HFD, and no significant differences were observed between the *ATP1A3* T807C and WT control mice (**Fig. 4C)**. Moreover, *ATP1A3* T807C mice showed a decreased percentage of body fat, accompanied by an increased percentage of body lean (**Fig. 4, D and E)**. The WAT of the whole epididymis was also significantly lighter in *ATP1A3* T807C mice than their WT counterpart mice (**fig. S4C)**. Measurement of metabolic parameters showed an increase of EE and VO_2_ in *ATP1A3* T807C compared to the WT controls (**Fig. 4, F and G)**. Furthermore, *ATP1A3* T807C mice showed enhanced glucose clearance during GTT (**Fig. 4H**), improved insulin sensitivity in ITT **(Fig. 4I**), and reduced HFD-induced liver steatosis compared to WT controls **(Fig. 4J**). Of note, inconsistent with what was observed in HLY72-treated mice (**Fig. 1, J and K**), *ATP1A3* T807C mice showed increased motion and RER in the metabolic cage studies compared to their WT counterparts (**fig. S5, D and E).** This may reflect a metabolic adaptation in *ATP1A3* T807C mice, wherein the significantly reduced body fat is compensated by increased carbohydrate utilization to support elevated energy expenditure (EE) and oxygen consumption (VO_2_). Meanwhile, the increased physical activity of *ATP1A3* T807C is consistent with the reported phenotype of *ATP1A3* I810N mice (I810N mutation reduces NKA α3 enzymatic activity by ∼40%) (*38*).

For the *ATP1A3* KO het mice, their body weight gains were also significantly decreased compared to their WT littermates (**Fig. 4K)**, with no difference in food intake (**Fig. 4L)**. *ATP1A3* KO het mice also showed a decreased percentage of body fat, an increased percentage of body lean, and a significantly lighter WAT compared to the WT control mice (**Fig. 4, M and N, and fig. S5F)**. Similar to what was observed in *ATP1A3* T807C mice, we also observed increased VO₂ (**Fig. 4, O)**, EE (**Fig. 4, P)**, and RER **(fig. S5G**) in *ATP1A3* KO het mice, along with significantly improved obesity-related complications, including enhanced glucose clearance during GTT (**Fig. 4Q**), improved insulin sensitivity in ITT (**Fig. 4R**), and reduced HFD-induced liver steatosis (**Fig. 4S**). *ATP1A3* KO het mice showed no difference in physical activity compared to their WT counterparts (**fig. S5H**).

Despite a similar ∼50% inhibition of NKA activity, *ATP1A3* T807C and *ATP1A3 KO* het mice are supposed to be different in their responses to HLY72 treatment. For this, we treated mice (WT, *ATP1A3* T807C and *ATP1A3* KO het) with a 60 kcal% HFD for 4 weeks followed by HLY72 treatment for another 3 weeks (**Fig. 4T)**. *ATP1A3* T807C mice gained significantly less body weight during HFD than their WT counterparts, and HLY72 induced a dramatic weight loss in WT but not in *ATP1A3* T807C mice (**Fig. 4, U and V**), further confirming that NKA α3 is the bona fide target of HLY72 *in vivo*. By contrast, *ATP1A3* KO het mice also gained significantly less weight compared to their WT littermates; however, HLY72 maintained its activity by further reducing weight gain in *ATP1A3 KO* het mice (**Fig. 4, W and X**).

Collectively, these results demonstrate that genetic inhibition of NKA α3, similar to chemical inhibition by HLY72, effectively protects mice from HFD-induced obesity and associated metabolic dysfunction.

## DISCUSSION

Our studies here revealed that NKA inhibitors with access to the brain are potential agents for counteracting obesity. Isoforms of NKA α subunits are highly homologous, and we do not think HLY72 has selectivity towards different α subunit isoforms. However, the anti-obesity effect of HLY72 is a specific outcome from targeting NKA α3, as reflected by the result that KI mice with the HLY72-binding site mutation in NKA α3 but not in the α1 gene abrogated HLY72’s activity (**Fig. 3, N to Q, and fig. S4, C to H)**. In line with the chemical inhibition of NKA α3 by HLY72, genetic inhibition of NKA α3 by the T807C mutation or heterozygous KO—both reducing NKA α3 activity by ∼50%—showed comparable protective effects against HFD-induced obesity and related metabolic problems such as hyperglycemia, insulin resistance, and hepatic steatosis, without affecting food intake (**Fig. 4, B to S)**. Inhibitors of NKA, such as CGs, have been used for centuries to treat congestive heart failure. In addition to CGs synthesized or isolated from plants, mammals including humans might also produce endogenous cardiac glycosides, mostly in the blood, adrenal gland, and hypothalamus (*39, 40*). However, our understanding of these endogenous cardiac glycosides is still quite incomplete. Our findings here implied that the *in vivo* regulation of NKA by some endogenous CGs, if it occurs, might play an important role in regulating energy homeostasis.

Our results showed that HLY72 could achieve potent anti-obesity efficacy without compromising energy intake at least within a certain dose range. Consistent with this, neither *ATP1A3* T807C mutation nor heterozygous KO exhibited any effect on food intake in mice. The unique property of HLY72 enables weight loss independent of reduction of food intake, thereby substantially reducing potential risks of complications typically associated with caloric restriction. On the other hand, our data showed that HLY72 induces statistically significant food intake reduction at higher doses (**Fig. 1D**). This appetite-suppressing effect of HLY72 is also mediated specifically by NKA α3, as demonstrated by the abolished response in NKA α3 T807C mutant mice (**fig. S4, C and D**). The mechanism behind the different effects of HLY72 on food intake remains enigmatic. NKA α3 exhibits high expression in multiple brain regions—including the cortex, basal ganglia, brainstem, cerebellum, hippocampus, and hypothalamus—with notable enrichment in GABAergic neurons. These regions are well-documented for their roles in feeding behavior and energy metabolism. HLY72 might exert divergent effects via NKA α3 in different brain regions, potentially due to receptor sensitivity variations across different neural circuits and cell populations. High doses of HLY72, due to the combined effect of its appetite-suppressing activity, might achieve a more pronounced anti-obesity effect; however, this may also lead to possible side effects. Understanding how HLY72 functions at different doses and their corresponding side effects is critical to establishing its therapeutic window and maximizing its anti-obesity potential. Intriguingly, during our study, mice treated with HLY72 exhibited significantly better conditions than those treated with digitoxin, with much less instances of drug-induced mortality even at high doses. Compared to digitoxin (> 24 hours), HLY72 has a very short half-life (∼ 5 min), as measured by the pharmacokinetic analysis (**Fig. 3I**). The rapid clearance of HLY72 *in vivo* might significantly reduce its toxicity, if it occurs. A recently reported anti-obesity BRINP2-related peptide (BRP) seems to support this hypothesis. The plasma half-life of BRP was quite short (less than 10 min), and it acts on the hypothalamus to induce appetite suppression while avoiding the typically associated side effects such as nausea or aversion (*41*). Therefore, development of drugs with transient target engagement might be a promising therapeutic strategy for safer anti-obesity treatment.

Our results here demonstrate that both chemical and genetic inhibition of NKA α3 effectively protect mice against HFD-induced adiposity and associated metabolic disorders, establishing NKA α3 as a promising therapeutic target for obesity management. Importantly, we provide a valuable anti-obesity strategy, which allows for obesity treatment and prevention without the need of altering energy intake.

---

**Supplementary Figure S1.**
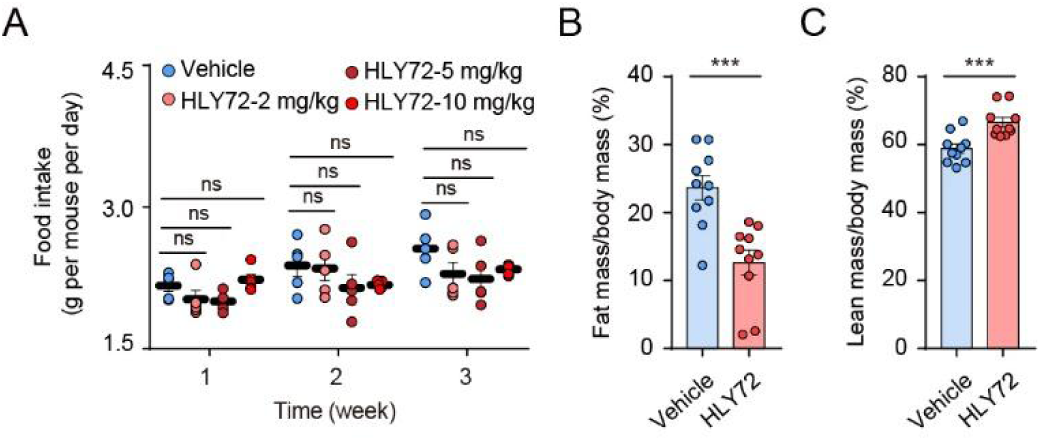
Effects of HLY72 on food intake and body fat. (**A**) Food intake of mice treated with vehicle or HLY72 at dosages of 2 mg/kg, 5 mg/kg, or 10 mg/kg. This figure is related to Figure 1F. (**B** and **C**) Percentage of fat mass (B), and percentage of lean mass (C). These two figures are related to Figure 1G and H respectively. Data are mean ± s.e.m. *p* values were determined by two-tailed un-paired student’s test in [(B) and (C)], two-way ANOVA in [(A)]. **\****p*<0.05, **\*\****p*<0.01, **\*\*\****p*<0.001. ns, not significant with *p*>0.05. Data points represent individual mice [(B) and (C)].

**Supplementary Figure S2.**
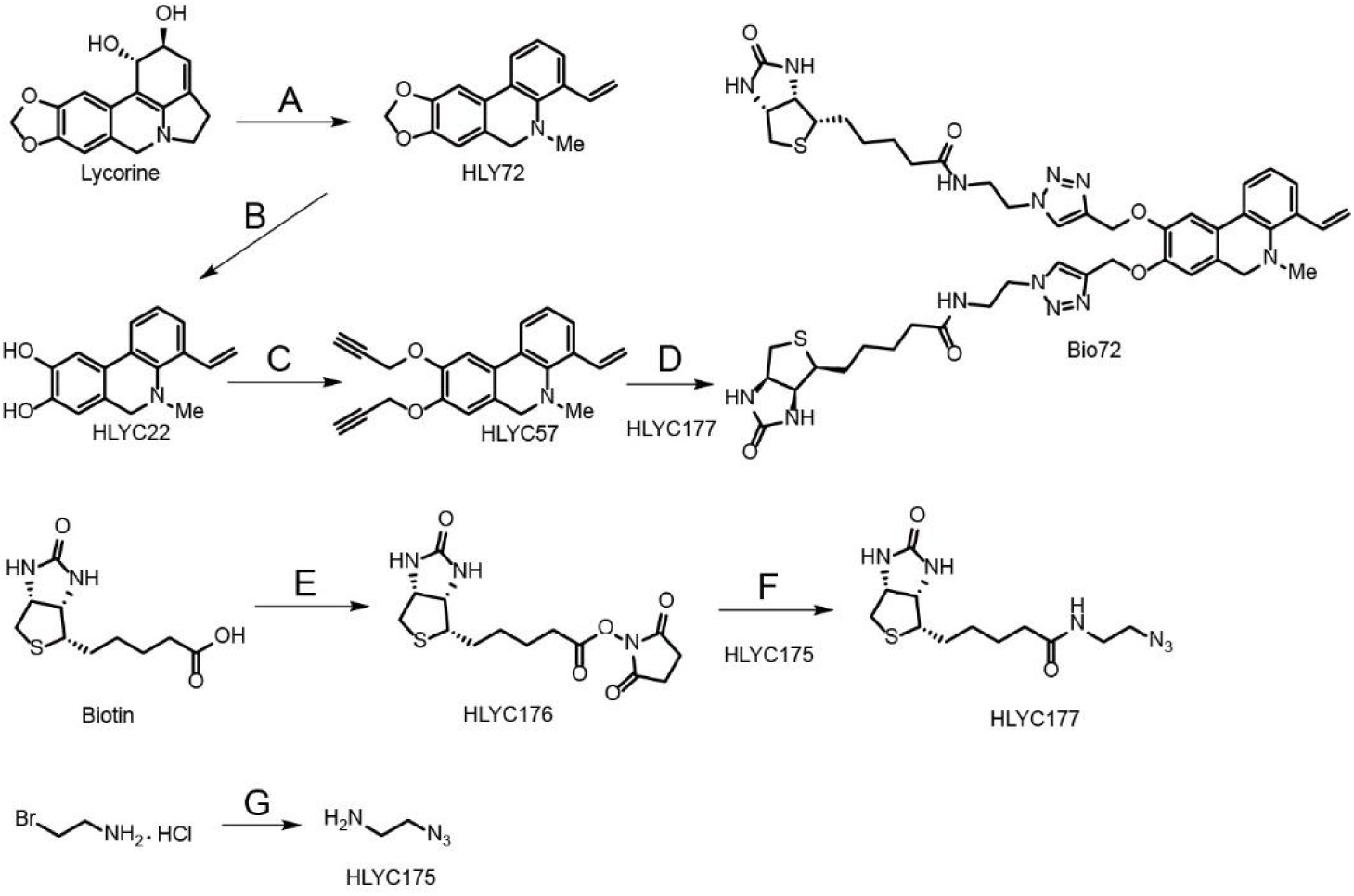
The syntheses of biotinylated HLY72 (Bio72). Reagents and Conditions. (**A**) 1.1. CH_3_l, room temperature, DMF, 12 hours; 1.2. KTB, TBA, 90℃, 4 hours, 85%. (**B**) Br_3_B, CH_2_Cl_2_, −78℃, 6 hours, 65%. (**C**) Propargyl bromide, K_2_CO_3_, THF, 75%. (**D**) HLYC177, Sodium ascorbate, CuSO_4_, TBA, 50 ℃ , 12 hours, 50% for Bio72. (**E**) N-Hydroxysuccinimide, DCC, pyridine, DMF, roomtemperature, 24 hours, 65%. (**F**) Et3N, DMF, room temperature, 24 hours, 70%. (**G**) Sodium azide, H2O, 75 ℃ , 21 hours, 82%. DMF: N, N-Dimethylformamide; KTB: Potassium tert-butoxide; TBA: Tert-Butyl Alcohol; DCC: Dicyclohexylcarbodiimide.

**Supplementary Figure S3.**
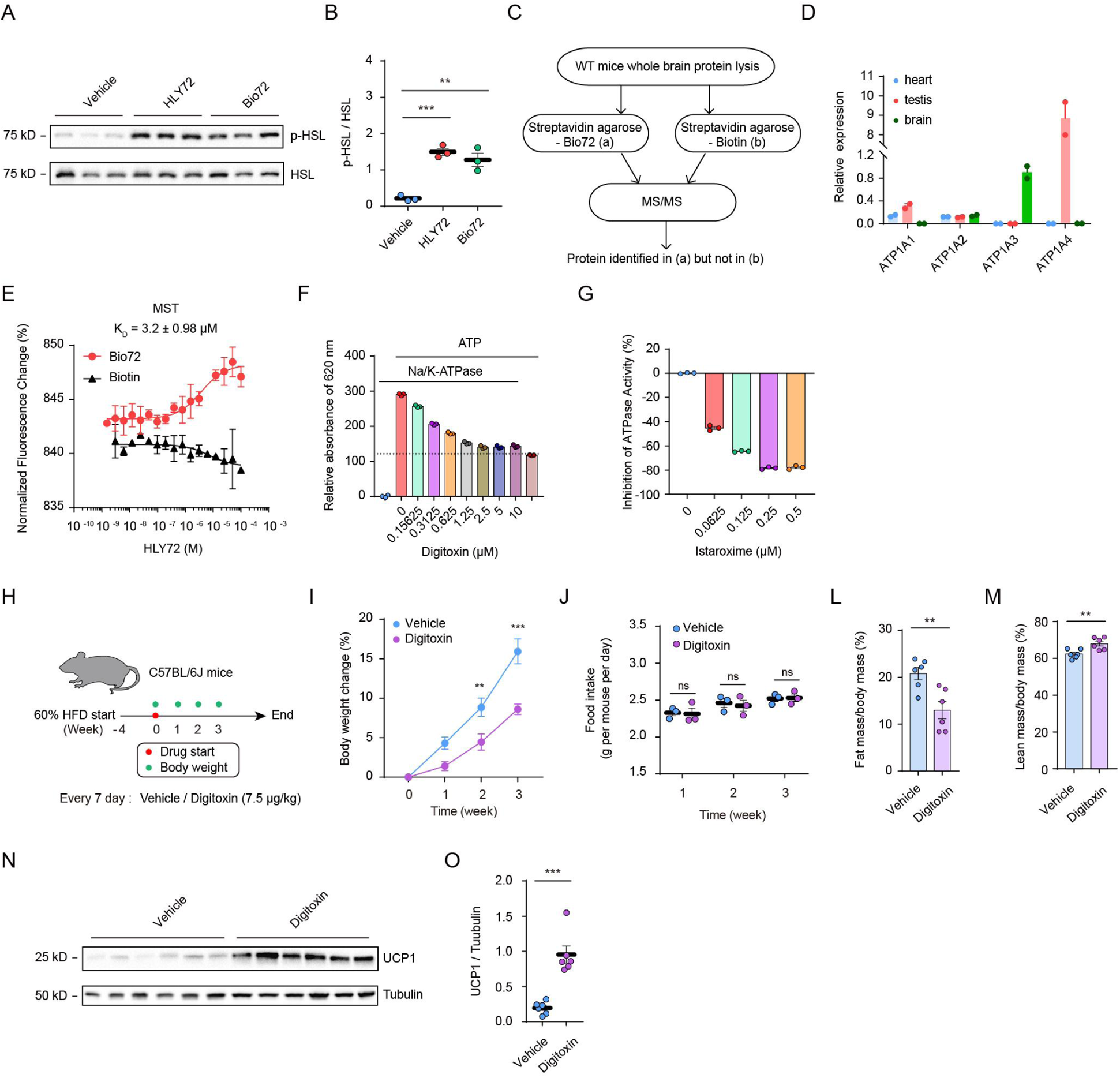
NKA α3 is identified as a target for HLY72. (**A** and **B**) p-HSL levels of 8-week-old male mice treated with HLY72 (10 mg/kg) and Bio72 (10 mg/kg) for 6 hours were examined by immunoblot analysis (A) and analyzed statistically (B). n=3. (**C**) Schematic scheme for target protein identification of HLY72 using MS. (**D**) Transcriptional levels of *ATP1A1*, *ATP1A2*, *ATP1A3* and *ATP1A4* genes in heart, testis and brain of mice. (**E**) The binding affinity of Bio72 with NKA were determined by using the MST assay. (**F**) Evaluation of ATPase activity of the purified NKA α3β1 protein by digitoxin. n=3. (**G**) Evaluation of the inhibitory activity of istaroxime against the purified NKA α3β1. n=3. (**H**) Experimental scheme for digitoxin (7.5 mg/kg) treatment in 8-week-old male mice. (**I** to **M**) Body weight (I), food intake (J), percentage of fat mass (L), and percentage of lean mass (M) were analyzed for mice administrated with digitoxin (7.5 mg/kg) for 3 weeks every 7 day. n=6. (**N** and **O**) UCP1 expression levels in BAT from mice treated with digitoxin (7.5 mg/kg) were examined by immunoblot analysis (N) and analyzed statistically (O). n=6. Data are mean ± s.e.m. *p* values were determined by two-tailed un-paired student’s test in [(L), (M) and (O)], one-way ANOVA in [(B)], two-way ANOVA in [(I) and (J)]. **\****p*<0.05, **\*\****p*<0.01, **\*\*\****p*<0.001. ns, not significant with *p*>0.05. Data points represent individual mice [(B), (L), (M) and (O].

**Supplementary Figure S4.**
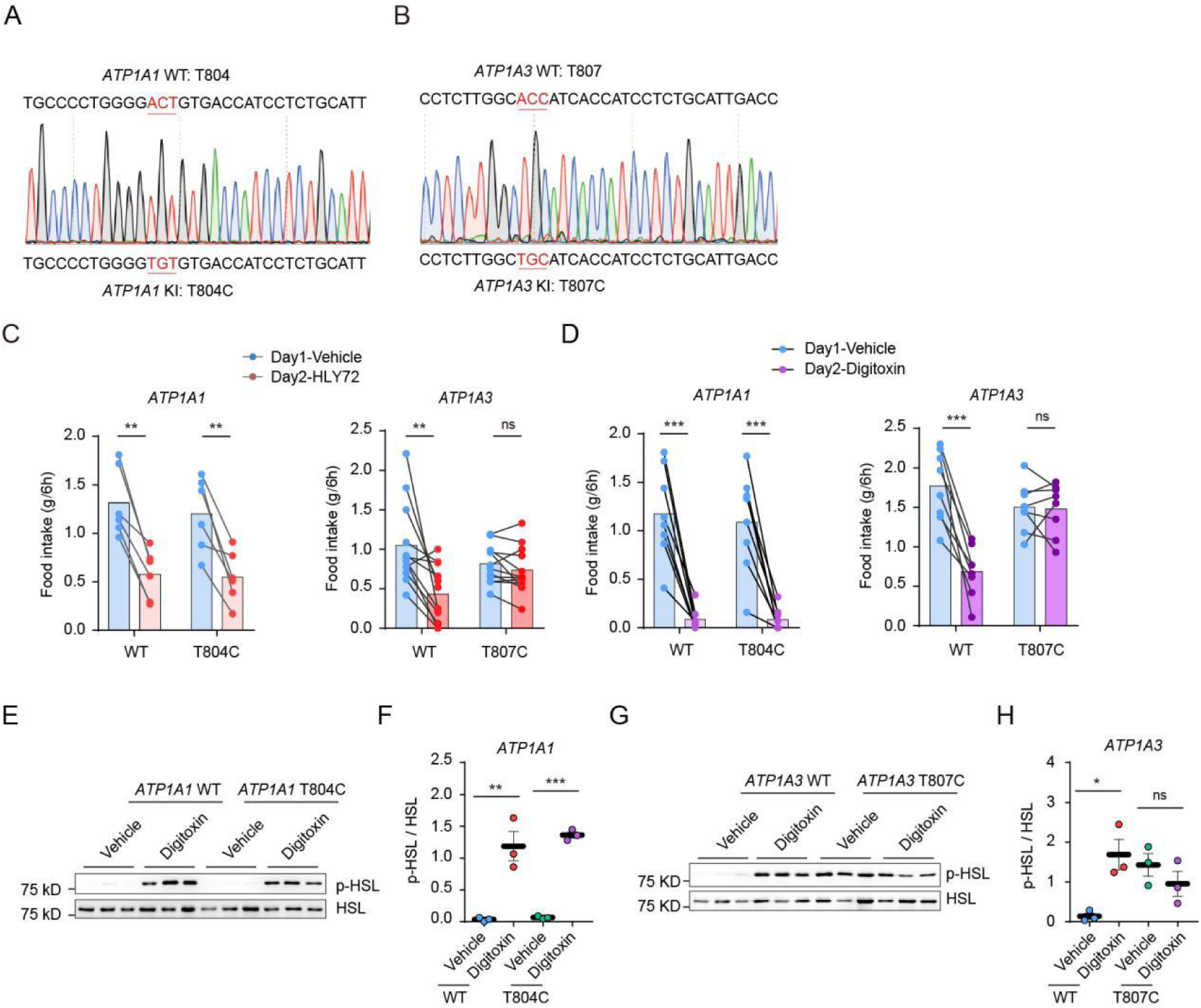
NKA α3 T807C mutation abolishes the effect of HLY72 in mice. (**A** and **B**) Sequencing confirmed that *ATP1A1* T804C and *ATP1A3* T807C mutant mice were successfully generated. (**C**) Food intake responses to 150 μg/kg digitoxin in 8-week -old male *ATP1A1* T804C and *ATP1A3* T807C mice, n=8. (**D**) Food intake responses to 20 mg/kg HLY72 in 8-week-old male *ATP1A1* T804C (n=6) and *ATP1A3* T807C mice (n=12). (**E** and **F**) p-HSL levels in total protein extracts of epididymal fats from 8-week-old male *ATP1A1* WT and T804C mice treated with digitoxin (150 mg/kg) were examined by immunoblot analysis (E), and analyzed statistically (F). n=3. (**G** and **H**) p-HSL levels in total protein extracts of epididymal fats from 8-week-old male *ATP1A3* WT and T807C mice treated with digitoxin (150 mg/kg) were examined by immunoblot analysis (G), and analyzed statistically (H). n=3. Data are mean ± s.e.m. *p* values were determined by two-tailed paired student’s test in [(C), (D)], two-tailed un-paired student’s test in [(F) and (H)]. **\****p*<0.05, **\*\****p*<0.01, **\*\*\****p*<0.001. ns, not significant with *p*>0.05. Data points represent individual mice [(C), (D), (F), (H)].

**Supplementary Figure S5.**
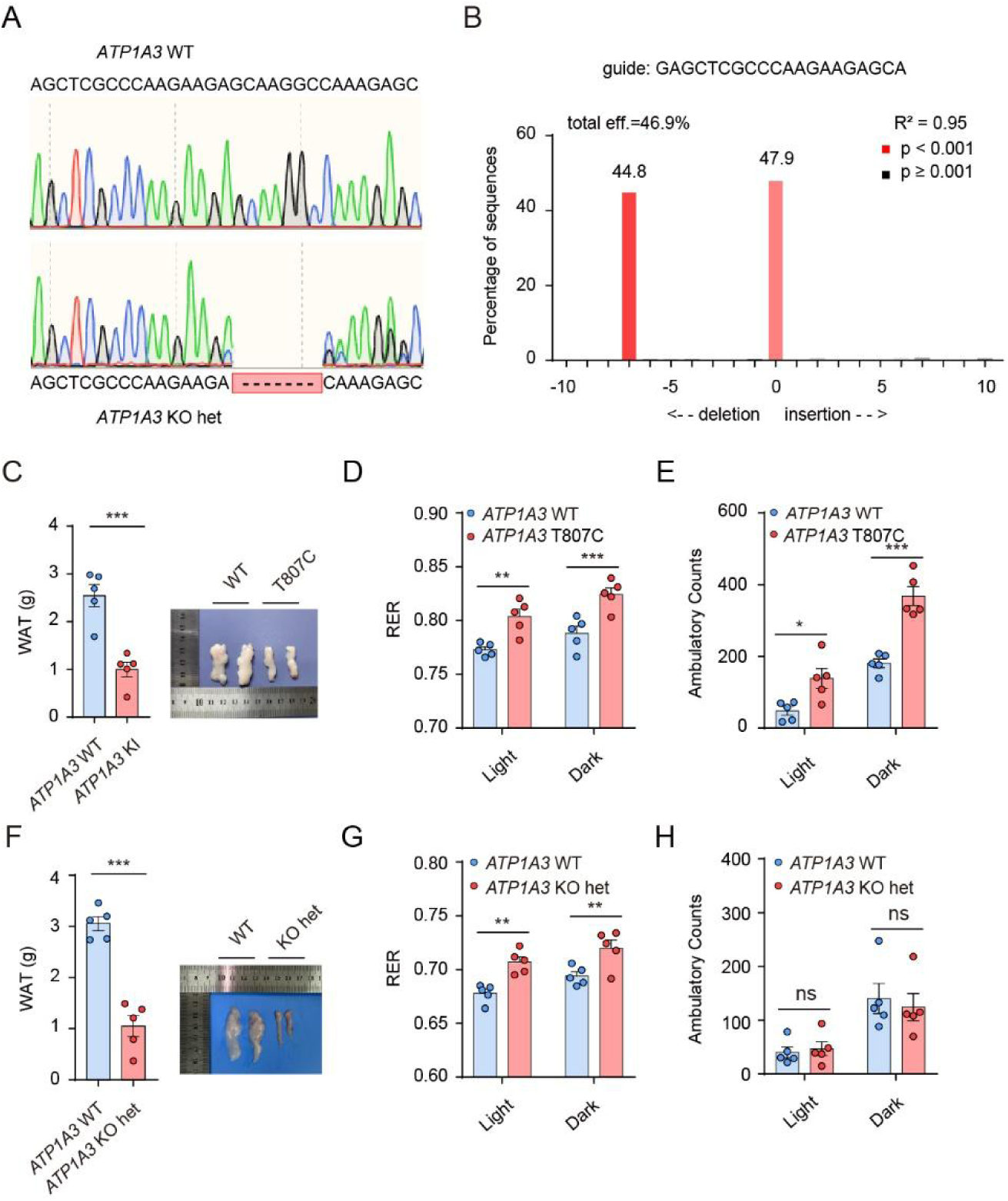
Generation of *ATP1A3* KO het mice and measurement of metabolic parameters of HFD-induced *ATP1A3* T807C and *ATP1A3* KO het mice. **(A** and **B**) Sequencing (A) and tracking of indels by decomposition (TIDE) assessment (B) to confirm that *ATP1A3* KO het mutant mice were successfully generated. **(C)** Representative figures of mouse eWAT in *ATP1A3* T807C *vs* WT mice. n=5. (**D** and **E**) RER (D) and ambulatory counts (E) of *ATP1A3* T807C *vs* WT mice. n=5. (**F**) Representative figures of mouse eWAT in *ATP1A3* KO het *vs* WT mice. n=5. (**G** and **H**) RER (G), and ambulatory counts (H) of *ATP1A3* KO het *vs* WT mice. n=5. Data are mean ± s.e.m. *p* values were determined by two-tailed un-paired student’s test in [(C) and (F)], two-way ANOVA in [(D), (E), (G) and (H)]. \**p*<0.05, \*\**p*<0.01, \*\*\**p*<0.001. ns, not significant with *p*>0.05. Data points represent individual mice [(C) to (H)].

